# Canine sense of quantity: evidence for numerical ratio-dependent activation in parietotemporal cortex

**DOI:** 10.1101/763300

**Authors:** Lauren S. Aulet, Veronica C. Chiu, Ashley Prichard, Mark Spivak, Stella F. Lourenco, Gregory S. Berns

## Abstract

The approximate number system, which supports the rapid estimation of quantity, emerges early in human development and is widespread across species. Neural evidence from both human and non-human primates suggests the parietal cortex as a primary locus of numerical estimation, but it is unclear whether the numerical competencies observed across non-primate species are subserved by similar neural mechanisms. Moreover, because studies with non-human animals typically involve extensive training, little is known about the spontaneous numerical capacities of non-human animals. To address these questions, we examined the neural underpinnings of number perception using awake canine functional magnetic resonance imaging. Dogs passively viewed dot arrays that varied in ratio and, critically, received no task-relevant training or exposure prior to testing. We found evidence of ratio-dependent activation, which is a key feature of the approximate number system, in canine parietotemporal cortex in the majority of dogs tested. This finding is suggestive of a neural mechanism for quantity perception that has been conserved across mammalian evolution.

## Introduction

Whether avoiding predators or foraging for food, it is evolutionarily imperative that animals can perceive and represent visual information regarding quantity. Extensive behavioral evidence suggests that non-human animals share with humans a sense of numerosity—in other words, a sensitivity to numerical information that does not rely on symbolic thought or education [1, 2]. This approximate number system (ANS), a system for rapidly assessing the approximate number of items present in an array, appears to be present both across the animal kingdom, as well as early in development, with even newborn infants possessing the remarkable ability to discriminate stimuli on the basis of numerosity [3].

Although many animals – including monkeys [4], fish [5], bees [6], and dogs [7, 8] – display behavioral sensitivity to numerosity, it is not clear whether the neural mechanism that underlies this ability is conserved across species. In humans, as well as non-human primates, evidence suggests that parietal cortex is the primary locus of this capacity [9, 10], although other cortical regions have also been implicated in this function [11, 12]. Recent work with birds, for example, suggests neural encoding of numerical information is localized in the endbrain of crows [13]. Given crows’ early divergence in phylogenetic history, it is not clear what the endbrain might be analogous to in mammals, or whether parietal mechanisms support numerical abilities in non-primates. To complicate the story, neural evidence for numerical abilities in non-human primates typically has required the animal to complete extensive training [7, 10]. Consequently, it is unclear what mechanisms underlie the capacities that these animals exhibit spontaneously [14, 15].

To address these questions, we used awake canine fMRI to assess numerical perceptual capacities in pet dogs. Because this methodology allows for assessment of number representations in the absence of number-specific training, and without relying on behavioral responses, we avoid common weaknesses in comparative numerical cognition (for further discussion of this issue, see [16]). In the present study, dogs passively viewed dot arrays that varied in numerical value. If dogs, like human and non-human primates, have a dedicated region of cortex for representing non-symbolic numerical quantity, then activation in this region should increase as the ratio between alternating dot arrays increases, despite constant cumulative surface area and variable element size [17]. That is, a number-sensitive region of cortex will exhibit greater activation when the numerical values of the stimuli presented are more dissimilar (e.g., 2 vs. 10 dots) than when numerical value is constant (e.g., 6 vs. 6 dots), consistent with Weber’s law [18].

## Methods

### Participants

Eleven awake, unrestrained dogs (see Table 1 for demographic information), were scanned in a Siemens 3T Trio MRI scanner. Prior to testing, all dogs completed a training program to be desensitized to the scanner environment through behavior shaping and positive reinforcement [19]. All dogs had previously participated in fMRI studies while viewing stimuli on a projection screen but had no prior training on numerical discrimination.

**Table 1.** Dogs’ demographic information

| **Dog** | **Breed** | **Sex** | **Age (in years)** |
| --- | --- | --- | --- |
| Bhubo | Boxer mix | M | 2 |
| Caylin | Border Collie | F | 10 |
| Daisy | Pitbull mix | F | 10 |
| Eddie | Labrador Golden mix | M | 7 |
| Kady | Labrador | F | 8 |
| Koda | Pitbull mix | F | 3 |
| Libby | Pitbull mix | F | 13 |
| Pearl | Golden retriever | F | 8 |
| Tallulah | Carolina dog | F | 6 |
| Truffles | Pointer mix | F | 13 |
| Zen | Labrador Golden mix | M | 9 |

### Stimuli

Stimuli were 75 dot arrays comprised of light gray dots on a black background (800 × 800 px). For each numerosity used (2, 4, 6, 8, and 10), stimuli varied in cumulative area (i.e., the total gray on each image). For each numerosity, cumulative area was either 10, 20, or 30% of the total stimulus. For each numerosity, 15 unique stimuli were used (5 stimuli per cumulative area value). In each stimulus, individual dot size varied up to 30%. Dot location varied randomly. Critically, these controls minimize the influence of non-numerical properties, in order to ensure that the results can be attributed to changes in numerical value [20, 21]. In accordance with current estimates of canine visual acuity (approximately 20/75; [22]), all stimuli were analyzed to ensure that the inter-dot distances were large enough for dogs to individuate.

### Block Design

During scanning, dogs passively viewed dot array stimuli presented on a screen placed in the rear of the scanner. Dogs were presented with alternating stimuli of 2 and 10 (1:5 ratio), 4 and 8 (1:2 ratio), or 6 and 6 (1:1 ratio) dots in a block fMRI design (see Figure 1). Stimuli were presented using PsychoPy software [23]. Each block contained 20 stimuli and lasted 10 seconds (see Supplemental Video). An experimenter standing in the rear of the scanner manually initiated each block to ensure that dogs were still in a suitable position within the scanner. Block onset times were recorded using an MRI-compatible button box. The inter-block intervals lasted approximately 10 seconds. Dogs were provided a food reward by their owner randomly throughout each run (always during the inter-block interval). Each run contained twenty blocks (randomized) and lasted approximately five minutes.

**Figure 1.**
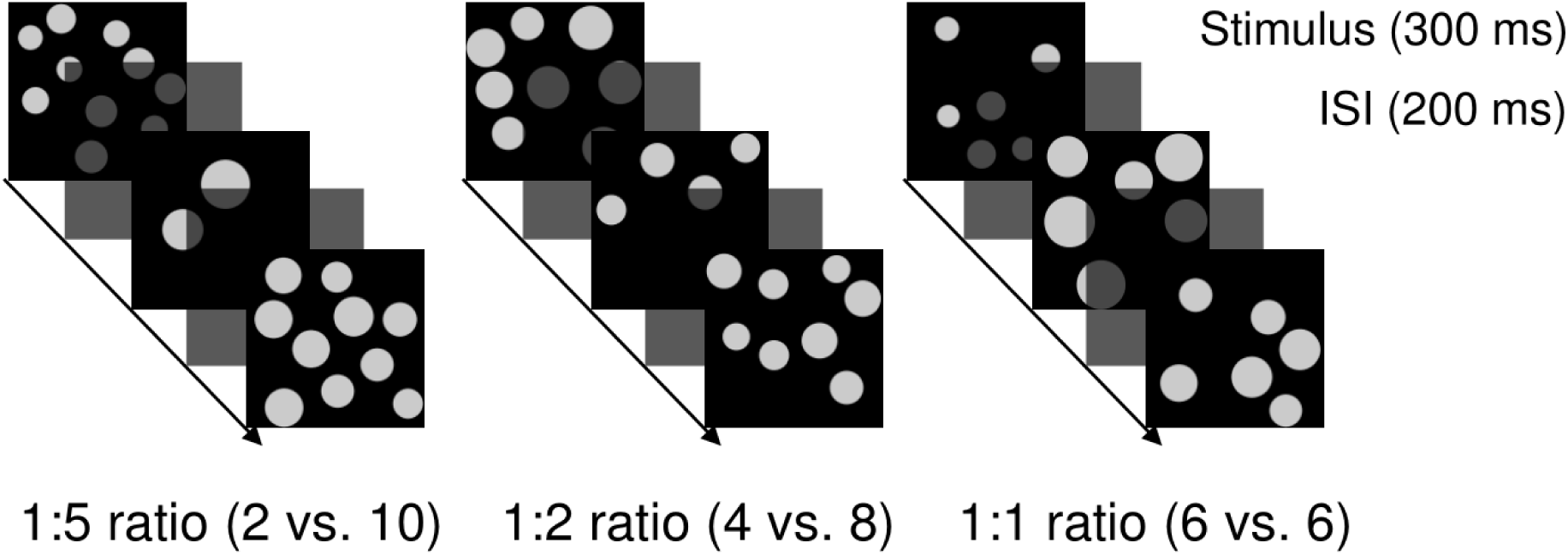
Block design of the present study.

### MRI Scan Acquisition

Dog training and fMRI protocol were consistent with the procedures previously used in awake dog fMRI studies [19, 24]. The scans were obtained using a Siemens 3T Trio MRI scanner. To obtain functional scans, a single-shot echo-planar imaging (EPI) sequence was used to acquire volumes of 22 sequential 2.5 mm slices with a 20% gap (TE = 25 ms, TR = 1260 ms, flip angle = 70 degrees, 64 × 64 matrix, 2.5 mm in-plane voxel size, FOV = 192 mm). For each individual, approximately 1300 to 2000 functional volumes were acquired over the course of two to five runs. For each dog, the total scan session lasted for a maximum of 40 minutes.

### Preprocessing

AFNI (NIH) was used for both preprocessing and statistical analysis. Preprocessing of the fMRI data included motion correction, censoring, and normalization using AFNI and its associated functions. Two-pass, six-parameter rigid-body motion correction was used based on a hand-selected reference volume for each dog that corresponded to the most representative position of the dog’s head within the magnet bore across runs. Aggressive censoring removed questionable volumes from the fMRI time sequence because dogs can move between trials and when consuming rewards. Censoring was performed with respect to both signal intensity and motion, in which voxels with greater than 3% signal change from the mean and volumes with more than 1 mm of scan-to-scan movement were flagged as spurious and censored from further analysis. Smoothing, normalization, and motion correction parameters were identical to those described in previous studies [25]. The Advanced Normalization Tools (ANTs) software [26] was used to spatially normalize the mean of the motion-corrected functional images to the individual dog’s structural image. To improve signal-to-noise ratio (SNR), the data were then spatially smoothed with a 4 mm gaussian kernel.

### Region of Interest (ROI) Analysis

Each dog served as its own control for cross-validation as we performed a four-fold split on the data. 75% of the data was used to localize the most likely number-selective region of interest (ROI) in each dog, while the remaining 25% was held out for independent validation. Because of the variability in response threshold across dogs, we used a customized approach that identified the most likely cluster of voxels correlated with number ratio in each dog. This was done by varying the voxel threshold of the statistical map for each dog so that one or two clusters remained that were 20-40 voxels in extent (see Supplemental Table 1). Once identified, the independent estimate from the holdout data was extracted from this ROI and submitted to a t-test across dogs.

For each dog, a General Linear Model (GLM) was estimated for each voxel using the 3dDeconvolve function in AFNI. Non-task regressors included the six motion regressors obtained from motion correction. Because neural activation was predicted to vary linearly with ratio, an amplitude-modulated response model was used in which a parametric modulator was assigned to each block based on the numerical ratio of the block (1, 2, or 5). This yielded two columns: a main effect of dot array stimuli and one modulated by the ratio of dots in a block. To allow for independent localization and test data, we further partitioned the design matrix into separate columns for either localization (75% of blocks) or testing (25% of blocks). We ensured that each test set had at least two instances of each condition represented in each run. To minimize any effect of fatigue or familiarization, we ensured that average onset times were matched (i.e., did not differ by > 5 seconds) across localization and test sets.

## Results

In eight of eleven dogs, we identified regions of cortex that exhibited increasing activation with numerical ratio in parietotemporal lobes according to a high-resolution canine brain atlas ([27]; see Figure 2A-C). Though there was variability in the specific location of each dog’s ROI, this is perhaps unsurprising given the different breeds in the current study [28] and evidence of non-parietal number-sensitive regions in the human literature [11].

**Figure 2.**
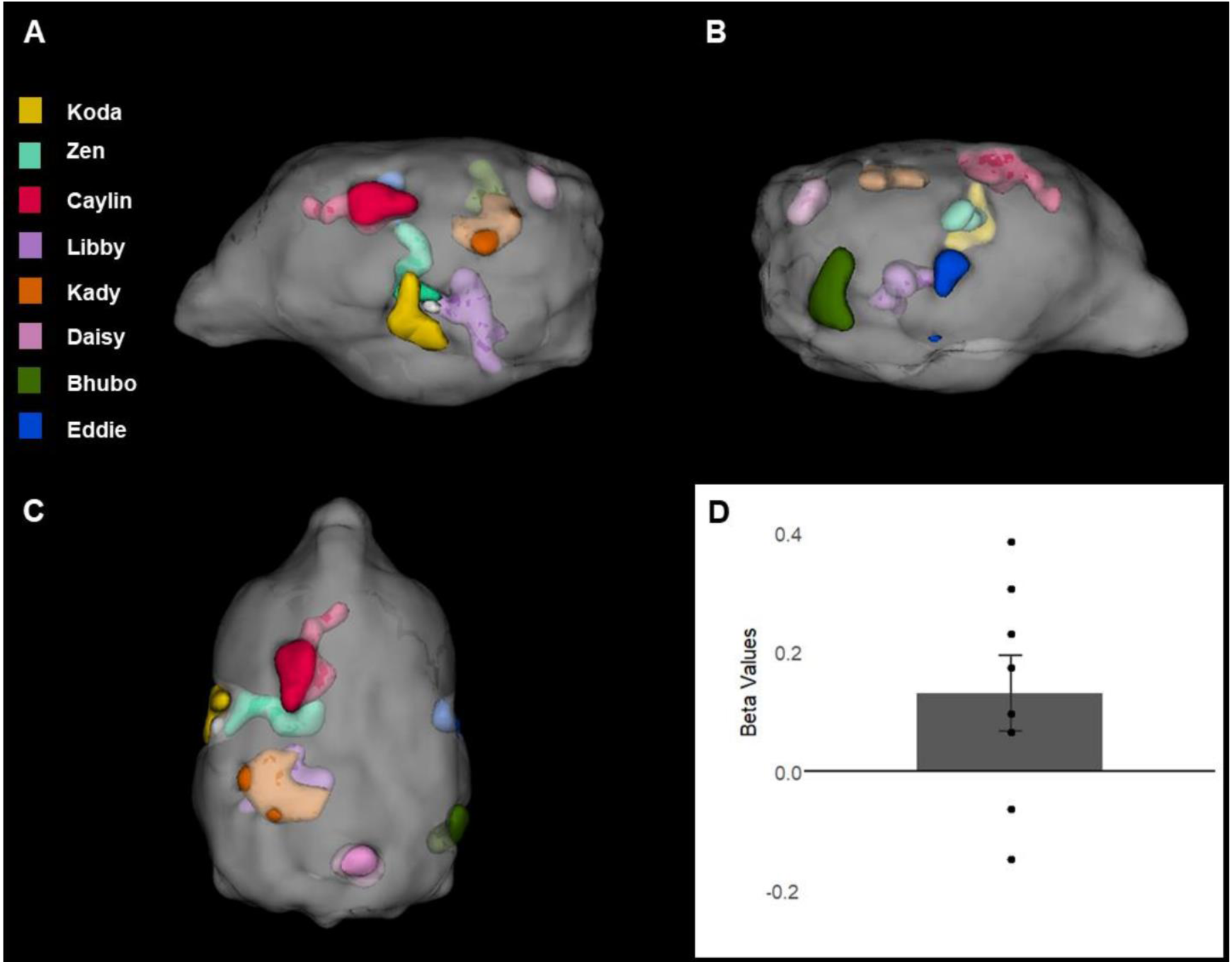
**(A-C)** Location of number-sensitive ROIs. Each color represents the ROI of one dog. For visualization and comparison of location, the ROIs have been spatially normalized and overlaid on a high-resolution dog brain atlas [27]. **(D)** Beta values from the number-sensitive ROIs in the held-out data for block ratio as a predictor of activation. Data points represent individual dogs. Error-bar is s.e.m.

Crucially, to assess whether dogs, like primates, have number-sensitive regions of cortex, we examined whether activation in the localized ROIs were ratio-dependent in the held-out data. We found that block ratio was significantly correlated with the level of activation in these regions [*t*(7) = 2.01, *p* = .042, one-sided; see Figure 2D], consistent with a ratio-dependent effect. These findings suggest that dogs have a visual sense of number and that this system is subserved by similar parietotemporal mechanisms to those in primates [10, 11].

## Discussion

In summary, we examined the neural underpinnings of the ANS in a novel mammalian species and, in the majority of dogs, found evidence of activation in parietotemporal regions that varied as a function of numerical ratio. Circumventing known weaknesses in the behavioral assessment of numerical abilities, especially in non-human animals [16], this work provides novel evidence that dogs do not require training in order to discriminate two-dimensional stimuli on the basis of non-symbolic number. Moreover, the present work suggests that dogs spontaneously discriminate dot arrays on the basis of numerical value even when arrays are equated for cumulative area and variable in individual element size. Though evidence from even a single animal provides an important proof of concept, the work here shows that the majority of dogs tested demonstrated spontaneous ratio-dependent neural activation, providing greater generalizability and stronger support for evolutionarily conserved neural mechanisms. Nonetheless, there are open questions about dogs’ ability to discriminate arrays on the basis of other ensemble properties such as average element size, which was variable in the current study, but still may have been discriminable. In naturalistic settings, however, such information is highly correlated with numerosity and likely to be used in combination with numerosity [29].

Consistent with recent evidence for emergent numerical capacities in computational models of object recognition [30], the present work provides evidence for neural representations of visual quantity, in the absence of explicit training. Moreover, the present study suggests that, like in humans, number perception occurs in parietotemporal cortex in dogs, a non-primate mammal. Taken together, our findings suggest that the ability to represent numerosity and the mechanisms supporting this system are deeply conserved over evolutionary time, perhaps due to a role in foraging or predation, and persists in a domesticated species.

## Acknowledgements

This work was supported by a National Institutes of Health (NIH) institutional training grant (T32 HD071845) to LSA, a Scholar Award from the John Merck Fund to SFL, and a grant from the Office of Naval Research (N00014-16-1-2276) to GSB. All views expressed are solely those of the authors. The authors would like to thank Raveena Chhibber for her contributions to this project.

## Author Contributions

LSA, SFL, and GSB designed the experiment. LSA, AP, VC, and GSB collected and analyzed the data. MS trained the dogs. LSA, VC, SFL, and GSB wrote the paper.

## Ethics Statement

This study was performed in accordance with the recommendations in the Guide for the Care and Use of Laboratory Animals of the National Institutes of Health. The study was approved by the Emory University IACUC (Protocols DAR-2002879-091817BA and DAR-4000079-ENTPR-A), and all owners gave written consent for their dog’s participation in the study.

## Supplemental Information

**Supplemental Table 1.** Number region size and threshold significance.

| **Dog** | **Voxels (*N*)** | ***p*** |
| --- | --- | --- |
| Bhubo | 20 | .05 |
| Caylin | 38 | .01 |
| Daisy | 13 | .05 |
| Eddie | 27 | .02 |
| Kady | 22 | .001 |
| Koda | 24 | .05 |
| Libby | 21 | .09 |
| Zen | 20 | .05 |

Dogs' number (ratio-dependent) ROI voxel size and *p* value for thresholds.

